# Floral resource-landscapes and pollinator-mediated interactions in plant communities

**DOI:** 10.1101/022533

**Authors:** Henning Nottebrock, Baptiste Schmid, Katharina Mayer, Céline Devaux, Karen J. Esler, Böhning-Gaese Katrin, Matthias Schleuning, Jörn Pagel, Frank M. Schurr

**Affiliations:** Institute of Landscape and Plant Ecology, University of Hohenheim, August-von-Hartmann-Str. 3, 70599 Stuttgart, Germany; Senckenberg Biodiversity and Climate Research Centre (BiK-F) and Senckenberg Gesellschaft für Naturforschung, Senckenberganlage 25, 60325 Frankfurt am Main, Germany; Department of Conservation Biology and Entomology and Centre for Invasion Biology, Stellenbosch University, Private Bag X1, Matieland 7602, South Africa; Institut des Sciences de l’Evolution, UMR 5554, Université Montpellier 2, Place Eugène Bataillon, 34095 Montpellier Cedex 05, France; Goethe University Frankfurt, Institute for Ecology, Evolution & Diversity, Max-von-Laue-Str.13, 60439 Frankfurt (Main), Germany

**Keywords:** bird foraging behaviour, community ecology, energetics of pollination, facilitation, indirect interactions among plants, intra- and interspecific competition, nectar sugar, plant-animal interactions, plant functional traits, species coexistence.

## Abstract

Plant communities provide floral resource-landscapes for pollinators. Yet, it is insufficiently understood how these landscapes shape pollinator-mediated interactions among multiple plant species. Here, we study how pollinators and the seed set of plants respond to the distribution of a floral resource (nectar sugar) in space and across plant species, inflorescences and flowering phenologies. In a global biodiversity hotspot, we quantified floral resource-landscapes on 27 sites of 4 ha comprising 127,993 shrubs of 19 species. Visitation rates of key bird pollinators strongly depended on the phenology of site-scale resource amounts. Seed set of focal plants increased with resources of conspecific neighbours and with site-scale resources, notably with heterospecific resources of lower quality (less sugar per inflorescence). Floral resources are thus a common currency determining how multiple plant species interact via pollinators. These interactions may alter conditions for species coexistence in plant communities and cause community-level Allee effects that promote extinction cascades.

## Introduction

Pollinators mediate indirect interactions between conspecific and heterospecific plants and can thus shape the dynamics of plant communities (Ghazoul 2005; Sargent & Ackerly 2008; Pauw 2013). Within plant populations, these pollinator-mediated interactions can be positive when neighbouring plants attract pollinators and increase visitation rates, or negative when plants compete for shared pollinators (Rathcke 1983; Ghazoul 2005). At the level of plant communities, generalist pollinators can mediate both competitive and facilitative interactions between plant species (Moeller 2004; Sargent & Ackerly 2008; Mitchell *et al*. 2009). These interspecific interactions depend on the foraging behaviour of pollinators in multi-species plants communities, and on whether interspecific pollen transfer reduces plant reproductive success (Waser 1978). Importantly, the relative magnitude of intra- and interspecific competition mediated by pollinators determines whether pollinators promote or hinder coexistence of plant species (Pauw 2013).

Energetic principles play a key role for pollinator-mediated interactions (Heinrich & Raven 1972; Heinrich 1975; Tomlinson *et al*. 2014): pollinators take up the energy provided by inflorescences (notably nectar) and partly use it for foraging movements that define the pollination services they deliver to plants. Consequently, spatial variation in the floral resource-landscape generated by a plant community should translate into spatial variation in pollinator foraging behaviour and pollinator-mediated interactions (Ghazoul 2005; Fig. 1a). Pollinator-mediated interactions also depend on flowering phenology because pollinators track temporal changes in resource-landscapes (Hegland *et al*. 2009; Fig. 1a). Despite these simple principles, pollinator-mediated interactions among plant species within communities can exhibit considerable complexity. This complexity arises from spatial and temporal variation in floral resources and from the partitioning of these resources among plant species and individual inflorescences (Fig. 1).

**Figure 1:**
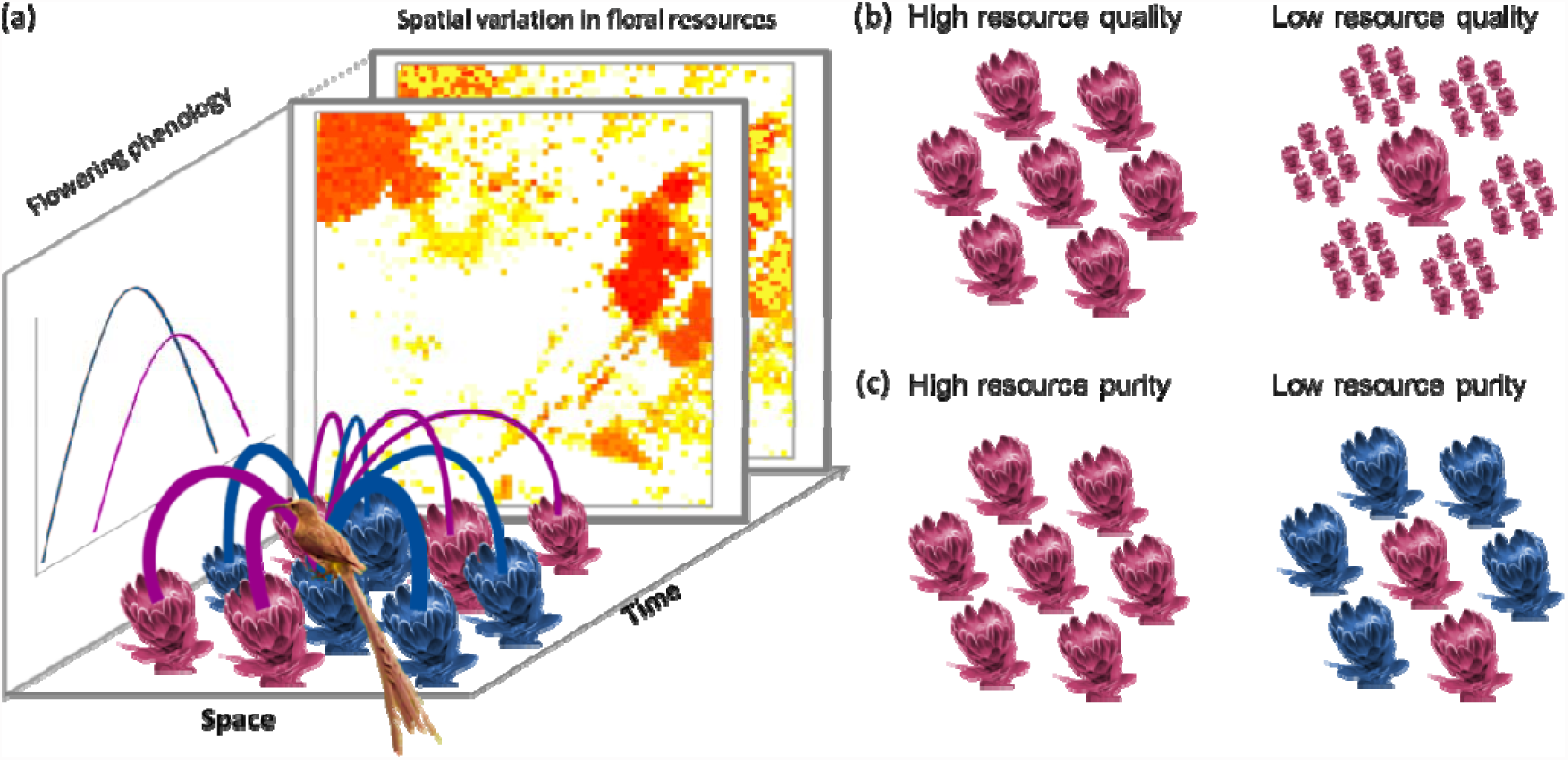
A conceptual framework for studying effects of floral resource-landscapes on pollinator-mediated interactions among plants. (a) Effects of spatial and phenological variation in floral resource amounts: the strength of pollinator-mediated interactions experienced by a focal inflorescence depends on the resource amount, spatial and phenological proximity of other inflorescences (interaction strength indicated by line widths). (b) Effects of floral resource quality: pollinator-mediated interactions depend on whether a given floral resource amount is split into a few high-quality inflorescences or into many low-quality inflorescences. In the example figures, the central inflorescence is either surrounded by inflorescences of equal quality (left) or lower quality (right). (c) Effects of floral resource purity: pollinator-mediated interactions depend on the proportion of conspecific floral resources. The example figures show cases of high purity (left) and low purity (right).

Pollinators can mediate interactions among plants at several spatial and temporal scales. Their small-scale foraging behavior affects interactions among inflorescences on the same plant (Goulson 2000; Devaux *et al*. 2014) while foraging movements determine interactions and pollen transfer among neighbouring plants (Seifan *et al*. 2014). At large spatial scales, pollinator abundance and pollination service respond to floral resource amounts provided by the entire community (Williams *et al*. 2012; Nottebrock *et al*. 2013). Importantly, the sign of pollinator-mediated interactions can change with spatial scale (Gunton & Kunin 2009). Overall, the intensity of pollinator-mediated interactions between two plants should decrease with the spatial and temporal distance between them (Heinrich & Raven 1972, Elzinga *et al*. 2007; Devaux & Lande 2009, Fig. 1a). Yet, even plants that do not flower simultaneously may interact via pollinators: early-flowering species can contribute to high pollinator densities that benefit late-flowering species (Riedinger *et al*. 2014).

In behavioural ecology, it is well established that the quality of resources in patches affects foraging decisions of animals. From the perspective of a foraging pollinator, an inflorescence is a food patch whose quality can be defined as the amount of floral resources available in a single visit (Pyke 1978). Hence, plant-pollinator interactions should not only depend on total resource amounts but also on whether these resources are split into a few high-quality inflorescences or into many low-quality inflorescences (Fig. 1b). Optimal foraging theory predicts that pollinators should respond to differences between the quality of a focal inflorescence and the quality of surrounding inflorescences: pollinators should prefer higher-quality inflorescences over lower-quality inflorescences (MacArthur & Pianka 1966) and they should spend more time visiting them (Charnov 1976; Pyke 1978). Higher-quality inflorescences can thus exert negative effects on pollinator visitation and reproductive success of surrounding plants with lower-quality inflorescences (Kandori *et al*. 2009). Conversely, higher-quality inflorescences could attract more pollinators, which then pollinate neighbouring plants with lower-quality inflorescences (Seifan *et al*. 2014). The net outcome of these opposite effects of higher-quality inflorescences on their surroundings remains unclear. Moreover, it is not obvious how quality differences between a focal inflorescence and other inflorescences should be evaluated, because the set of available inflorescences depends on the spatial scale at which pollinators take their foraging decision, which is generally poorly known (Ghazoul 2005).

Pollinator-mediated interactions between a focal plant and the surrounding floral resources can also be affected by the ‘purity’ of these resources, defined as the proportion of floral resources contributed by conspecifics (Fig. 1c, Ghazoul 2005). Positive effects of purity on pollinator efficiency and plant reproductive success result from increased intraspecific pollen transfer and reduced stigma clogging by incompatible heterospecific pollen (Waser 1978; Shore & Barrett 1984). Additionally, purity may increase reproductive success via positive effects on pollinator visitation (Ghazoul 2005) because pollinators preferentially visit common plant species or because they sequentially visit inflorescences of the same species (Chittka and Thomson 2001). On the other hand, purity can reduce plant reproductive success if competition for pollinators is more intense among conspecifics than among heterospecifics (Pauw 2013). Furthermore, heterospecifics can increase pollinator visitation if different plant species with temporally staggered flowering phenologies facilitate each other via the maintenance of high pollinator densities (Riedinger *et al*. 2014). Hence, the purity of floral resources can have either positive or negative effects on plant reproductive success and the balance between these effects most probably varies with the spatial and temporal scales at which floral resource purity is considered.

The spatial distribution, phenology, quality and purity of floral resource-landscapes are thus expected to strongly shape pollinator-mediated interactions among plants. Previous studies considered these aspects individually, demonstrated their relevance for plant-pollinator interactions but also yielded seemingly conflicting results (e.g. Kunin 1997; Ghazoul 2005, Gunton & Kunin 2009; Williams *et al*. 2012; Carvalheiro *et al*. 2014; Feldman & McGill 2014). We argue that progress in understanding the effects of floral resources on pollination requires an integrative approach that quantifies the aforementioned aspects of floral resource-landscapes and analyses their relative importance for pollinator behaviour and plant reproductive success (Fig. 1). Here, we develop such an approach and apply it to 27 plant communities from the South African Fynbos biome, a global biodiversity hotspot (Myers *et al*. 2000). Our objectives are to (1) quantify how floral resource-landscapes vary in space, time, quality and purity, and (2) determine the relevance of these aspects of floral resource-landscapes for pollinator visitation and seed set. We show that floral resource-landscapes explain pollinator-mediated interactions within and among plant species. Importantly, the multi-scale impacts of floral resources on plant communities can alter conditions for species coexistence and can cause community-level Allee effects that promote extinction cascades.

## Material and Methods

### Study system and study design

We studied shrub communities dominated by the species-rich genus *Protea* that has high ecological and economic importance in the Fynbos biome (Schurr *et al*. 2012) and is well suited for studying plant-pollinator interactions. *Protea* species frequently dominate the overstorey of Fynbos shrublands and provide copious amounts of nectar accumulated at the base of their inflorescences (flowerheads) (Collins & Rebelo 1987). These inflorescences bear many individual florets, each of which contains a single ovule and can thus produce a single seed (Rebelo 2001). To set seed, *Protea* species require pollinator visits to inflorescences and many species are strongly dependent on pollination by nectarivorous birds, notably Cape sugarbirds (*Promerops cafer*) and orange-breasted sunbirds (*Anthobaphes violacea*, Schmid *et al*. 2015). Since inflorescences (referred to as cones after flowering) are the functional unit of plant-pollinator interactions in our study system, we measured standing nectar sugar crops, pollinator visitation and seed set at the level of inflorescences.

Making use of the high beta-diversity of *Protea* meta-communities, we selected 27 study sites that vary in species composition and density of *Protea* (Fig. 2a). Each site consisted of a 200×200 m^2^ plot with a core zone of 120×120 m^2^ surrounded by a 40 m wide buffer zone (Fig. 2b). To analyse the effects of floral resource-landscapes on pollinator-mediated interactions at these sites, we (1) generated fine-scale maps of all overstorey *Protea* individuals, (2) quantified sugar amount per inflorescence and phenological variation in the number of flowering inflorescences to predict floral resource-landscapes (Fig. 2d), (3) measured both visitation rates of key bird pollinators and seed set at the inflorescence level for a further subset of plants, and (4) and ran statistical analyses that quantify how pollinator visitation and seed set are shaped by floral resources at the plant, neighbourhood and site scale, and by the phenology, quality and purity of these floral resources.

**Figure 2:**
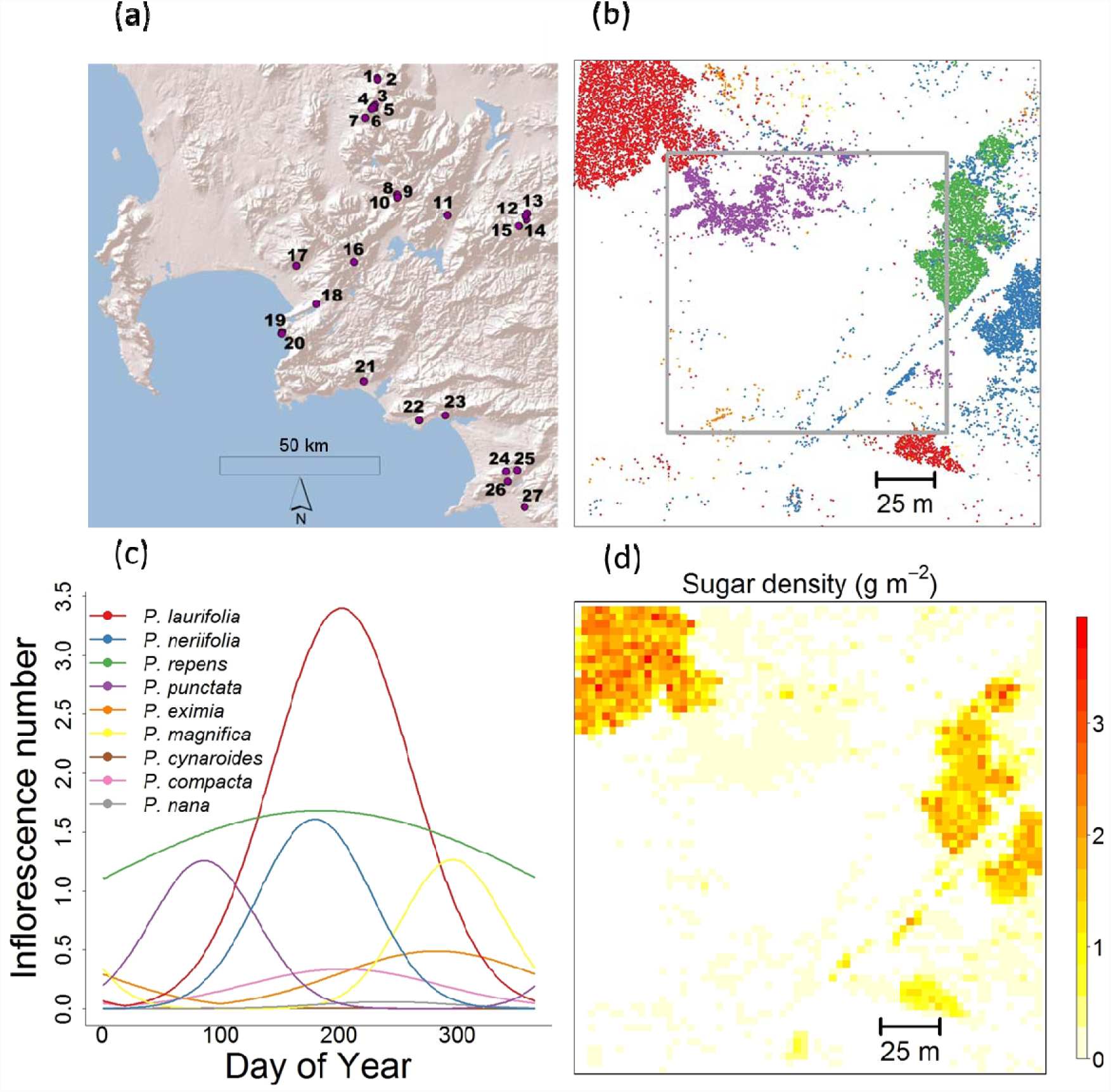
Quantifying the spatiotemporal dynamics of floral resource-landscapes. (a) Location of 27 study sites in the Fynbos biome, South Africa. (b) Map of 16,948 shrub individuals on study site 4 with colours indicating different Protea species (see legend in (c)). (c) Flowering phenologies of the nine Protea species on this site (shown as the number of flowering inflorescences of a median-sized plant). (d) Spatial distribution of nectar sugar on the site predicted for a given day (4 July).

### Fine-scale mapping

We mapped all overstorey *Protea* plants on the study sites using differential GPS (Trimble GeoXH; median accuracy 20 cm) and recorded their size (canopy height) and species identity. In very dense monospecific stands (>6 individuals per 2 m^2^), we mapped the stand outline, recorded plant density and then simulated plant locations within the stand according to a complete spatial random distribution with the observed density. The sizes of these simulated plants were drawn from a stand-specific gamma distribution estimated by a maximum likelihood fit to the sizes of 30 plants measured per stand. In total, the fine-scale maps comprise 127,993 individuals of 19 *Protea* species, with 318 to 48,602 individuals per species, 83 to 37,253 individuals per site, and 3 to 9 species per site.

### Trait-based prediction of floral resource-landscapes

We monitored individual flowering phenologies for a subsample of 6,943 plants (51 to 1245 plants per species) by counting flowering inflorescences at up to three visits during the flowering seasons in 2011 (March to December) or 2012 (March to August). For a subsample of 850 plants in the core zones, (4 to 80 plants per species) we harvested two inflorescences, measured their size and the proportion of open florets, and extracted their nectar by centrifugation (Armstrong & Paton 1990). We measured nectar volume with microsyringes (0.05 mL precision) and nectar concentration with a hand refractometer (Bellingham and Stanley, reading range: 0-50 Brix). Nectar concentration in Brix was then converted into grams of sugar per litre and multiplied with nectar volume to obtain sugar amount per inflorescence.

To predict floral resource landscapes, we fitted trait-based models of sugar amount per inflorescence and number of inflorescences per plant. As predictors for these trait-based models, we measured inflorescence size, cone mass, specific leaf area (SLA), and trunk length from the ground to the first branch for a subsample of 2,580 plants in the core zone (25 to 502 plants per species). Additionally, the models included resprouting ability as a species-level trait (Rebelo 2001). The model for inflorescence number also included a date-derived covariate to describe species-specific flowering phenologies. With these trait-based models we then predicted phenological variation in inflorescence number, sugar amount per inflorescence and their product, sugar amount per plant, for all 127,993 mapped plants (for details see Appendix S1 in Supporting Information).

From these spatially explicit predictions, we derived the amount, quality and purity of floral resources at the neighbourhood and site scales. At the neighbourhood scale (within 40 m radius around each focal plant), we calculated sugar amounts using a neighbourhood index that accounts for the decline of neighbour effects with distance d from the focal plant (Uriarte et al. 2010): we summed the sugar amounts of all neighbours within 40 m weighted by 1/(1+*d*). At the site scale, we calculated the total sugar amount of all plants on the site (in g/ha). At both the neighbourhood and site scales, we also calculated purity and resource quality. Purity was calculated as the proportion of the sugar amount at the respective scale that is contributed by conspecifics of the focal plant. As a relative measure of resource quality at the neighbourhood and site scale, we subtracted the focal plant’s sugar per inflorescence from the mean sugar per inflorescence at the respective scale.

Phenology was treated differently when characterizing floral resource-landscapes for analyses of pollinator visits and seed set, respectively (see below). For pollinator visits, we considered floral resource-landscapes at the respective day of observation. In contrast, seed set integrates over the entire flowering period of an inflorescence and seed set analyses thus included temporally averaged resource variables that were weighted by the phenology of the focal plant (Appendix S1).

### Pollinator observations and seed set measurements

Pollinator visitation and seed set were measured on plants located within the core zones of the study sites. On up to three visits per site we counted legitimate inflorescence visits by nectarivorous Cape sugarbirds (*Promerops cafer*) and orange-breasted sunbirds (*Anthobaphes violacea*). We recorded the number of inflorescences probed by birds for 1,333 plants (1 to 346 plants per species) during 45 min sessions in the morning (8am – 10am, up to 10 plant-level observations per session). We only considered legitimate probing events, in which birds had contact with stigmas and thus potentially transferred pollen.

Seed set was measured for 1,717 plants (22 to 378 plants per species) by counting the number of fertile seeds (W_fertile_) in up to five randomly harvested mature cones (Nottebrock *et al*. 2013). The seeds were cross-cut and then probed with a needle to identify fertile seeds containing a soft endosperm. Pre-dispersal seed predation rate was estimated as the proportion of the cross-sectional cone area consumed by predators. The total number of ovules per plant that could potentially set seed was calculated as W_potential_= (1- π_p_) A_C_ / A_S_, where π_*p*_ is the estimated predation rate, A_C_ and A_S_ are the cross-sectional areas of cones and seeds (A_C_ was measured for each cone, A_S_ was determined as the mean of up to 50 seeds per population).

### Analysing effects of floral resource-landscapes on pollinator-mediated interactions

To analyse how pollinator visits and seed set respond to different aspects of floral resource-landscapes, we used generalised linear mixed models (GLMMs, package lme4, Bates *et al*. 2014) in R 3.1.1 (R Core Team 2013). We used Poisson errors for the number of pollinator visitations and binomial errors for seed set expressed as the ratio of fertile seeds to potential seeds (W_fertile_/W_potential_). The model for pollinator visitation controlled for the number of visible inflorescences per plant (included as an offset) in order to describe pollinator visitation rate per inflorescence.

As explanatory variables, the models for both response variables included measures of floral resources at three spatial scales: the number of inflorescences and sugar per inflorescence at the focal plant scale, and sugar amount at the neighbourhood and site scales. To describe how resource purity and quality modify the effects of sugar amount at the neighbourhood and site scale, we included interactions of purity and quality with sugar amounts at the respective scale. We did not include main effects of purity and quality since this would imply that purity and quality play a role when sugar amounts are zero. To facilitate the interpretation of purity effects, we used impurity (1-purity), which is zero for a purely conspecific neighbourhood. Hence, the main effects of sugar amounts describe effects of ‘pure’ resource-landscapes in which all sugar is provided by conspecifics. By adding the impurity-interaction term to the corresponding main effect of sugar amount, one obtains the effect of sugar provided exclusively by heterospecifics with identical resource quality. The further addition of the quality-interaction term describes the effect of sugar provided by heterospecifics with higher resource quality.

Analyses of both pollinator visitation and seed set corrected for focal plant size and the seed set analysis additionally controlled for direct plant-plant interactions (such as competition for nutrients) by including the density of con- and heterospecific neighbours (using again the 1/(1+*d*) distance-weighting index). Lastly, we accounted for random variation in space, time and among species: for pollinator visits we included random effects of plant species and observation session (which encompasses site and day effects) and for seed set we included random effects of plant species and site.

To quantify the relevance of different aspects of floral resource-landscapes for pollinator visitation and seed set, we calculated the AIC difference between the full models (see above) and control models without the respective aspect. Control models for different spatial scales were obtained by dropping all resource variables at the respective scale, whereas control models for resource quality and purity omitted the respective interaction terms. In the control model for phenology, we replaced all phenology-weighted resource variables by the respective annual mean.

Finally, we examined the relationship between seed set (response variable) and pollinator visitation (explanatory variable) for the 279 plants for which both data were available. We used a binomial GLMM with a fixed effect of visitation per inflorescence and random effects of species identity and site. Note that pollinator observations were conducted on single dates within the flowering season, but not necessarily at the plant’s peak flowering time. Pollinator visitation rates that were observed close to a plant’s peak flowering time can be expected to be more representative for the entire flowering period and thus more closely related to seed set than visitation rates observed towards the limits of the plant’s flowering period. We therefore weighted each data point by exp((-Δt^2^)/σ), where Δt is the time difference between the pollinator observation and the plant’s peak flowering time and σ is the standard deviation of the plant’s flowering phenology (Appendix S1).

## Results

### Spatiotemporal variation of floral resource-landscapes

Trait-based models of flowering phenology and sugar amount per inflorescence quantify the spatiotemporal dynamics of floral resource-landscapes in the 27 study communities (Fig. 2, Video S1). At the plant scale, sugar per inflorescence varied between 0.01 g and 1.94 g, and the annual maximum of co-flowering inflorescences per plant varied between 0 and 44. The 19 study species showed considerable differences in flowering phenology: their peak flowering time varied from March to October and they ranged from temporally-peaked to year-round flowering (Fig. 2c, Table S1). We calculated the average floral resource-landscape experienced by a flowering inflorescence by integrating sugar amounts and inflorescences over these flowering phenologies (see Appendix S1). At the site scale, this phenology-integrated sugar amount was on average 388.9 g/ha (95% interquantile range: 11.1 – 1414.9 g/ha) with a mean purity of 52% (0 – 99%). The mean sugar amount of co-flowering inflorescences on the same site differed from an inflorescence’s own sugar amount by an average quality difference of +0.008 g (-0.7 – +0.8 g). The summed sugar amount in the neighbourhood of flowering inflorescences (weighted by 1/(1+*d*)) was on average 18.3 g (0.4 – 103.3 g) with a mean purity of 63% (0 – 100%) and a mean quality difference of -0.003 g (- 0.6 – +0.7 g).

### Effects of floral resource-landscapes on pollinator visits and seed set

The spatial structure, quality, purity and phenology of floral resource-landscapes were of different relevance for pollinator visitation and seed set (Fig. 3). For pollinator visitation, the relevance of floral resources at different spatial scales increased from the plant over the neighbourhood to the site scale (Fig. 3a). Visitation rates depended strongly on the phenology of floral resources, and to a lesser extent on resource quality, but resource purity was of minor relevance for pollinator visitation (Fig. 3a). In contrast, seed set was mostly driven by floral resources at the neighbourhood scale (Fig. 3b). Moreover, seed set was strongly affected by the purity of floral resource-landscapes, whereas resource quality had intermediate relevance and phenology had relatively minor relevance for seed set (Fig. 3b).

**Figure 3:**
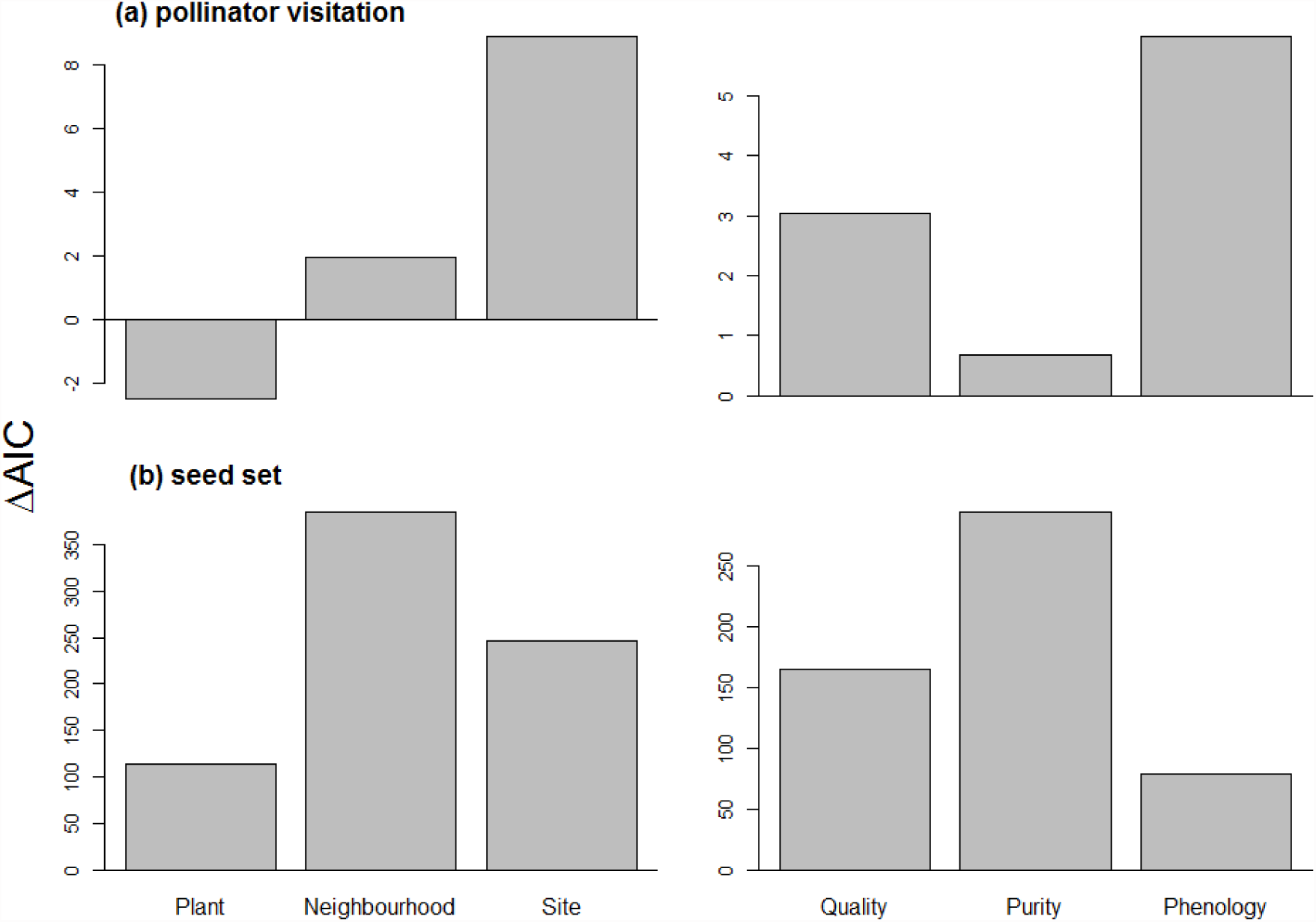
Relevance of different aspects of floral resource-landscapes for (a) pollinator visitation per inflorescence and (b) seed set per inflorescence. The left panels show the relevance of floral resources at three spatial scales, the right panels show the relevance of floral resource quality, purity, and phenology. The relevance of a given aspect of resource-landscapes is measured as the AIC difference difference between a control model model without the respective aspect and the full model (a positive value indicates better performance of the full model).

Significant effects of floral resource-landscapes on pollinator visitation were only found at the neighbourhood and site scales, where the main effects of sugar amount were modified by interactions with resource quality (Fig. 4a). Pollinator visitation increased with sugar amount at the neighbourhood scale if neighbouring inflorescences had higher resource quality than the focal inflorescence (positive quality-resource interaction, (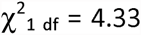, P < 0.05, Fig. 4a). Site-scale sugar amounts had a strong negative effect on pollinator visitation, which was particularly pronounced if site-scale sugar amounts were composed of higher-quality inflorescences (negative quality-resource interaction, (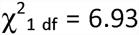, P < 0.01, Fig. 4a). In contrast, the purity of floral resources did not alter the effect of sugar amount on pollinator visitation at either scale (P > 0.05).

**Figure 4:**
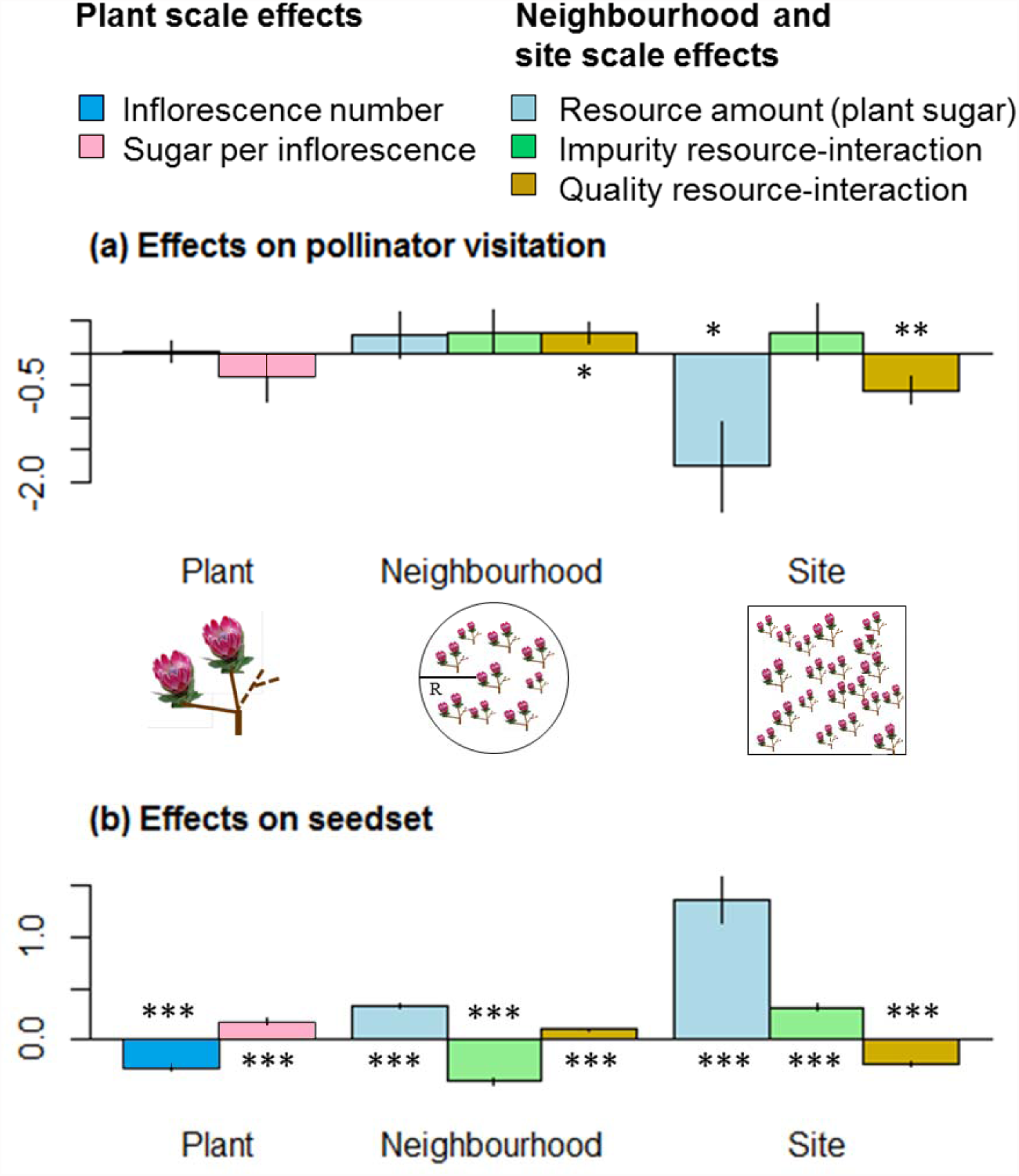
Effects of floral resource-landscapes at the plant, neighbourhood and site scale on (a) pollinator visitation and (b) seed set per inflorescence. Bars indicate standardized regression coefficients, whiskers the corresponding standard errors and stars the significance of effects (*: p < 0.05, **: p < 0.01, ***: p < 0.001). At the plant scale, bars show the effect of inflorescence number (dark blue) and sugar amount per inflorescence (pink). At the neighbourhood and site scale, light blue bars show main effects of sugar amount, green bars show interactions between impurity (proportion of heterospecific sugar) and sugar amount, and brown bars show interactions between relative resource quality (difference in sugar per inflorescence) and sugar amount. Light blue bars at the neighbourhood and site scale thus represent effects of purely conspecific sugar amounts, the addition of the corresponding green bars yields the effect of heterospecific sugar amounts with identical quality, and the addition of the corresponding brown bars shows how resource effects are altered for heterospecifics with higher resource quality.

Seed set showed significant responses to all aspects of floral resource-landscapes at all spatial scales (Fig. 4b). At the plant scale, seed set increased with sugar amount per inflorescence (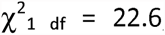, P < 0.001, Fig. 4b) and decreased with the number of inflorescences on the focal plant (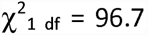, P < 0.001, Fig. 4b). At the neighbourhood scale, seed set increased with floral resource amounts consisting entirely of conspecific sugar (positive main effect of neighbour sugar amount), but slightly decreased with resource amounts consisting entirely of heterospecific sugar (the positive main effect of neighbour sugar amount was outweighed by the negative impurity-resource interaction, (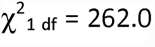, P< 0.001, Fig. 4b). This negative effect was particularly pronounced if neighbouring inflorescences had lower quality than the focal inflorescence (positive quality-resource interaction, (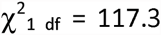, P < 0.001, Fig. 4b). While floral resource neighbourhoods had either positive or negative effects on seed set (depending on resource purity and quality), the effects of neighbour plant density were consistently negative. The negative intraspecific density dependence of seed set was stronger than the negative interspecific density dependence (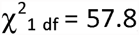,P < 0.001). This negative effect of conspecific density was almost exactly compensated by the positive effect of conspecific sugar amounts (standardized regression coefficients for conspecific density and sugar amount were -0.33 and +0.33, respectively, Fig. 4b). At the site scale, we found a strong positive effect of sugar amounts, which was more positive if site-scale sugar resources were provided by heterospecific plants (positive impurity-resource interaction, (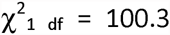, P < 0.001) and by lower-quality inflorescences (negative quality-resource interaction, (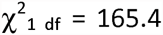, P < 0.001, Fig. 4b). A positive relationship between pollinator visitation and seed set was found for the 279 focal plants on which we had measured both variables. The seed set of these plants showed a logistic response to pollinator visitation rate (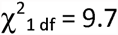, P < 0.01).

## Discussion

The high-resolution description of floral resource-landscapes for 27 plant communities enabled us to quantify how floral resources (nectar sugar amounts) vary in space and time, and how their partitioning among plant species and inflorescences causes differences in resource purity and quality. The relevance of these aspects of floral resource-landscapes differed between pollinator visitation and seed set: pollinator visitation largely depended on site-scale floral resources, whereas seed set was determined jointly by floral resources at the plant, neighbourhood and site scales (Figs. 3 and 4). Here we discuss the mechanisms causing these floral resource effects and their consequences for the dynamics of plant communities.

### Floral resource effects on pollination and seed set

Floral resource amounts at the site scale had a strong negative effect on pollinator visitation per inflorescence but a strong positive effect on seed set (Fig. 4). While the negative response of pollinator visitation may seem surprising, it can be explained by the behaviour of bird pollinators. On the same study sites, bird pollinator abundance increases less than proportional with site-scale resources (B. Schmid, *personal communication*), possibly due to territoriality of bird pollinators. This negative effect does, however, not propagate into seed set (Fig. 4b). The opposite response of seed set to site-scale floral resources could result from saturation of stigmas at relatively low levels of pollinator visits, above which more visits do not translate into higher seed set. We observed such a saturating effect in the logistic relationship between seed set and pollinator visitation. Importantly, any interpretation of the differential responses of pollinator visitation and seed set to site-scale resource amounts must consider the different temporal scales at which pollinator-mediated interactions act: competition for pollination results mainly from the behavioural response of pollinators to instantaneous resource offers, whereas facilitation mainly results from the numerical response of pollinators to long-term resource availability (Gahzoul 2005; Riedinger *et al*. 2014). Facilitative effects caused by increased pollinator abundance thus likely dominate the positive effect of phenology-integrated resource variables on seed set. In contrast, pollinator visitation was negatively related to floral resource availability on the same day, which likely results from short-term competition for pollinator visits.

The purity of floral resources had weak effects on visitation (Figs. 3a and 4a), which is consistent with the finding that the bird pollinators of our study species are generalists that visit all available study species (B. Schmid, *personal communication*). In contrast, seed set increased with the purity of floral resources in the neighbourhood and decreased with the number of inflorescences on the focal plant (Fig. 4b), which is expected if seed set is limited by the availability of outcrossed conspecific pollen. The larger importance of phenology for pollinator visitation rather than seed set could arise because pollinator visitation depends on instantaneous resource-landscapes at the day of pollinator observation, whereas seed set integrates over phenological variation throughout the season. These different temporal scales could also explain why the positive effect of the site-scale floral resources on seed set increased with impurity (Fig. 4b) so that heterospecific floral resources had a stronger facilitative effect than conspecific resources. The flowering phenologies of our study species are displaced (Fig. 2b), which should reduce interspecific competition for shared pollinators (Devaux & Lande 2009). On the other hand, facilitative effects via the maintenance of high pollinator populations through the season are enhanced by the staggering of flowering phenologies among species (Moeller 2004; Riedinger *et al*. 2014). Overall, the balance between competitive and facilitative effects on pollination visitation and seed set can thus be more positive for heterospecific than for conspecific floral resources.

The resource quality (sugar per inflorescence) of focal plants had a positive effect on their seed set (Fig. 4). Moreover, pollinator visitation and seed set of plants with lower-quality resources benefitted from higher-quality neighbours, which suggests that these neighbours attract pollinators and exert a ‘magnet effect’ (Moeller 2004; Seifan et al. 2014). In contrast, it is disadvantageous for a plant to offer resources of lower quality than the site-scale average. This possibly arises because the large-scale foraging decisions of pollinators induce site-scale competition for pollination.

### Floral resources and plant community dynamics

The role of floral resources and pollinator-mediated interactions for the dynamics of plant communities has received increasing attention in recent years (Sargent & Ackerly 2008; Pauw 2013; Greenspoon & M’Gonigle 2013). We found that both intra- and interspecific floral resources at the site scale have strong positive effects on plant reproductive success. Previously, Nottebrock *et al*. (2013) found positive effects of large-scale community density on seed set and lifetime fecundity of *Protea* repens. The present study of 19 *Protea* species in 27 communities suggests that such community-level Allee effects are a general feature of *Protea* communities and that they are mediated by floral resources. Community-level Allee effects can have profound consequences for plant population and community dynamics: decreased floral resources of certain plant species can increase the extinction risk of other plant species, thus increasing the susceptibility of communities to extinction cascades (Colwell *et al*. 2012).

Our findings also have interesting implications for species coexistence and the structure of diverse plant communities. We found that seed set in *Protea* communities is affected by negative direct effects of plant density and by predominantly positive effects of floral resources (Fig. 4b). The direct density effects reveal that intraspecific density-dependence is more negative than interspecific density-dependence, which should cause rare species to experience less competition than common species and should therefore stabilize coexistence (Chesson 2000). These stabilizing density effects are, however, counteracted by pollinator-mediated effects at the neighbourhood scale: conspecific floral resources increase seed set whereas heterospecific resources have much weaker effects (Fig. 4b). These resource-based effects thus tend to neutralize intraspecific competition while leaving interspecific competition unaffected. Hence, an individual plant immigrating into a neighbourhood dominated by another species will have strongly reduced seed set compared to a member of the dominant species. This ‘priority effect’ should promote the formation of monospecific stands (M’Gonigle & Greenspoon 2014) that are a prominent feature of *Protea* communities (cf. Fig. 2a). The emergence of such monospecific stands reduces neighbourhood-scale coexistence but can facilitate larger-scale coexistence. This is because stable stand boundaries slow down large-scale competitive exclusion which led M’Gonigle & Greenspoon (2014) to state that it ‘stabilizes coexistence’. In the classification of Chesson (2000), however, this effect is equalizing (reducing fitness differences between species) rather than stabilizing (favouring rare species). In contrast, the positive effects of site-scale floral resources on seed set (Fig. 4b) are stabilizing sensu Chesson (2000): site-scale facilitation is stronger between than within species, which favours species that are rare at the site scale.

Our results suggest that pollinator-mediated interactions contribute to the formation of monospecific stands, but cause interspecific facilitation across stand boundaries, which stabilizes site-scale coexistence. These effects can help to explain the typical spatial structure of plant communities in the biodiversity hotspot studied here, which differs from other megadiverse systems (such as tropical forests) through the existence of monospecific stands at small scales, but high species richness at larger scales and thus high beta-diversity (Goldblatt & Manning 2002). Such multi-scale impacts of pollinator-mediated interactions on plant communities are not fully covered by existing single-scale theories (Sargent & Ackerly 2008; Pauw 2013; Greenspoon & M’Gonigle 2013).

### Conclusions and Outlook

This study shows that floral resources are a common ‘interaction currency’ (Kissling *et al*. 2012) that determines how multiple plant species interact via their shared generalist pollinators. It identifies inflorescence number and sugar amount per inflorescence as key quantities that convert the spatial structure and phenology of individual plant species into the spatiotemporal dynamics, purity and quality of this common currency at the community level. Pollinator visitation and seed set respond to these multiple aspects of the floral resource currency, with potentially important consequences for the dynamics and coexistence of plant species within communities. The identification of such interaction currencies is crucial for both developing a more general understanding of community dynamics and predicting community dynamics in changing environments (McGill *et al*. 2006; Kissling *et al*. 2012). It is timely to test whether resource-landscapes play similar roles in other pollination systems and for other types of generalized trophic interactions, such as plant-herbivore and plant-frugivore networks.

## Acknowledgements

This paper is dedicated to the memory of the late Brummer Olivier without whom this study would not have been possible. For discussion and input we are grateful to Anton Pauw, Phoebe Barnard and Tony Rebelo. We also thank a large number of field and lab assistants. We were able to accomplish our study thanks to the support of private land-owners (Giel von Deventes, Flower Valley Conservation Trust; Mathia, Nayna and Walter Heidehof, Gansbaai; Bairie and Peter Gibson, Macially High Noon Farm G. and S. Moskovitz) and nature reserves (Grootbos, Fernkloof, Helderberg, Hottentots-Holland, Jonaskop, Limietberg, Mont-Rochelle). Field work was conducted under Cape Nature permit AAA005-00213-0028. This work was supported by the German Research Foundation (grant numbers SCHU 2259/3-1, SCHL 1934/1-1). This is publication number ISEM X-X.

## Supporting Information

Additional Supporting Information may be downloaded via the online version of this article at Wiley Online Library (www.ecologyletters.com).

As a service to our authors and readers, this journal provides supporting information supplied by the authors. Such materials are peer-reviewed and may be re-organized for online delivery, but are not copy-edited or typeset. Technical support issues arising from supporting information (other than missing files) should be addressed to the authors.

